# Maintenance of taste receptor cell presynaptic sites requires gustatory nerve fibers

**DOI:** 10.1101/2024.07.28.604832

**Authors:** Shannon M. Landon, Emily Holder, Amber Ng, Ryan Wood, Eduardo Gutierrez Kuri, Laura Pinto, Saima Humayun, Lindsey J. Macpherson

**Affiliations:** The University of Texas at San Antonio, Department of Neuroscience, Developmental and Regenerative Biology, One UTSA Circle, San Antonio, TX, USA; Brain Health Consortium, The University of Texas at San Antonio.

**Keywords:** Taste, Gustatory, Chemosensory, Synapse, Synaptogenesis, Plasticity

## Abstract

The turnover and re-establishment of peripheral taste synapses is vital to maintain connectivity between the primary taste receptor cells and the gustatory neurons which relay taste information from the tongue to the brain. Despite the importance of neuron-taste cell reconnection, mechanisms governing synapse assembly and the specificity of synaptic connections is largely unknown. Here we use the expression of presynaptic proteins, CALHM1 and Bassoon, to probe whether nerve fiber connectivity is an initiating factor for the recruitment of presynaptic machinery in different populations of taste cells. Under homeostatic conditions, the vast majority (>90%) of presynaptic sites are directly adjacent to nerve fibers. In the days immediately following gustatory nerve transection and complete denervation, Bassoon and CALHM1 puncta are markedly reduced. This suggests that nerve fiber innervation is crucial for the recruitment and maintenance of presynaptic sites. In support of this, we find that expression of *Bassoon* and *Calhm1* mRNA transcripts are significantly reduced after denervation. During nerve fiber regeneration into the taste bud, presynaptic sites begin to replenish, but are not as frequently connected to nerve fibers as intact controls (∼50% compared to >90%). This suggests that gustatory neuron proximity, rather than direct contact, likely drives taste receptor cells to express and aggregate presynaptic proteins at the cell membrane. Together, these data support the idea that trophic factors secreted by gustatory nerve fibers prompt taste receptor cells to produce presynaptic specializations at the cell membrane, which in turn may guide neurons to form mature synapses. These findings provide new insights into the mechanisms driving synaptogenesis and synaptic plasticity within the rapidly changing taste bud environment.

## Introduction

The mammalian taste bud contains a collection of mature and developing taste receptor cell (TRC) populations, as well as the peripheral axons of gustatory neurons. Together, they detect and then relay taste signals from the tongue to the brain. TRCs detect one of the 5 major taste modalities: sweet, bitter, salty, sour, and umami. TRCs that respond to sweet, bitter, and umami are classified as Type II cells and have unique presynaptic structures whereby the neurotransmitter, ATP, is released through CALHM1/3 channels (Taruno et al., 2013; Ma et al., 2018; Romanov et al., 2018; Kashio et al., 2019). These channels are restricted to points of contact with afferent nerve fibers, allowing for precise, non-vesicular neurotransmission (Romanov et al., 2018). Type III cells, which detect sour and ionic stimuli, release serotonin through classical vesicular exocytosis (Yang et al., 2000; Huang et al., 2008; Liman and Kinnamon, 2021). While numerous synaptic markers have been evaluated for type III cells, Bassoon, a scaffolding protein in presynaptic active zones, has proven to be the only consistently reliable marker for identifying type III cell presynaptic sites (Yang et al., 2000, 2007, 2020; Kohno et al., 2006; Asano-Miyoshi et al., 2009; Kotani et al., 2013; Ikuta and Hamada, 2022).

Taste information is primarily relayed from TRCs via ATP signaling, activating the purinergic P2X2/P2X3 receptors specific to gustatory neurons within the taste bud (Bo et al., 1999; Finger et al., 2005; Eddy et al., 2009; Huang et al., 2011; Vandenbeuch et al., 2015). There is evidence showing serotonin plays a small role in taste signaling (Larson et al., 2015), however, the activation of P2X2/P2X3 receptors necessitates the transduction of every taste modality, including sour (Finger et al., 2005; Vandenbeuch et al., 2015; Flammer et al., 2024). To relay these taste signals, the peripheral axons of gustatory neurons associate closely with TRCs, often making more than one synaptic contact (Wilson et al., 2021). The synaptic contacts made between TRCs and gustatory neurons do not last long, as TRCs have a finite lifespan of approximately 10 days (Beidler and Smallman, 1965). Older TRCs die off and are replaced by newly born TRCs, a frequent event commonly referred to as TRC turnover. Innervating gustatory neurons must, then, re-connect with appropriate TRCs on a regular basis. Therefore, the formation and removal of synaptic connections in the taste bud is quite common. This continual synapse turnover is critical for the integrity of taste sensation and taste related behaviors, however, mechanisms governing synapse formation in the taste bud are entirely unstudied.

It is likely that processes in both taste cells and gustatory neurons coordinate to organize synapse formation in the taste bud. Here, we probe the connectivity between gustatory neurons and TRC presynaptic proteins to interrogate the role of gustatory nerve fiber innervation and contact in synaptogenesis within the taste bud. Our goal is to determine whether TRCs rely on gustatory neuron innervation for the formation of presynaptic sites and whether neuronal contact triggers the assembly of presynaptic sites on TRCs. To accomplish this, we employed the use of presynaptic markers, Bassoon and CALHM1, and gustatory neuron marker, P2X2, to observe gustatory nerve fiber innervation and contact frequencies with presynaptic sites on TRCs in normal and dernervated conditions. We found that presynaptic sites are highly colocalized with gustatory nerve fibers, which then effectively vanish following denervation. When gustatory nerves re-innervate the taste bud, presynaptic sites return on TRCs, though they are contacted far less frequently than intact controls. Thus, we propose that TRCs depend entirely on the close proximity of gustatory neurons to produce presynaptic sites, independent of direct contact. This finding provides a basis for mechanisms driving the continual re-establishment of synapses within the taste bud.

## Results

### Presynaptic specializations are strongly associated with nerve fiber connectivity

Are the presynaptic sites on TRCs always occupied by gustatory neurons? Answering this question will illuminate the extent to which TRCs rely on gustatory neuron innervation to form and maintain presynaptic sites. To determine this, we used immunostaining of presynaptic sites on type II (CALHM1) and type III cells (Bassoon), observed as fluorescent puncta within the taste buds. Additionally, we stained for gustatory neurons (using the purinergic receptor P2X2) to examine exactly how many presynaptic sites (puncta) are contacted by gustatory neurons under homeostatic conditions. We analyzed fungiform taste buds found at the front of the tongue and circumvallate taste buds at the back of the tongue. We determined that the average number of Bassoon puncta is 3.41 in fungiform taste buds and 7.47 in circumvallate taste buds (Figure 1a-c). These Bassoon puncta are contacted by nerve fibers at a frequency of 89% for fungiform and 95% for circumvallate taste buds (Figure 1d). On average, there are more CALHM1 puncta in taste buds than Bassoon, with fungiform taste buds containing an average of 15.8 puncta and circumvallate taste buds containing an average of 26.5 puncta (Figure 1e-g). Our observation of a higher puncta-per-bud count for Calhm1 compared to Bassoon is expected due to the greater abundance of type II cells relative to type III cells in each taste bud (Ogata and Ohtubo, 2020; Yang et al., 2020; Wilson et al., 2021). Furthermore, CALHM1 puncta are contacted by nerve fibers 91% of the time for both fungiform and circumvallate taste buds (Figure 1h). Both Bassoon and CALHM1 boast a high percentage of puncta contacted by nerve fibers, which supports the idea that nerve fiber innervation and contact are important for accumulation of presynaptic sites in TRCs. However, there still is a small percentage (5-11%) of Bassoon and CALHM1 puncta that are not contacted, indicating that neuronal contact is not absolutely necessary for TRCs to produce presynaptic sites.

**Figure 1:**
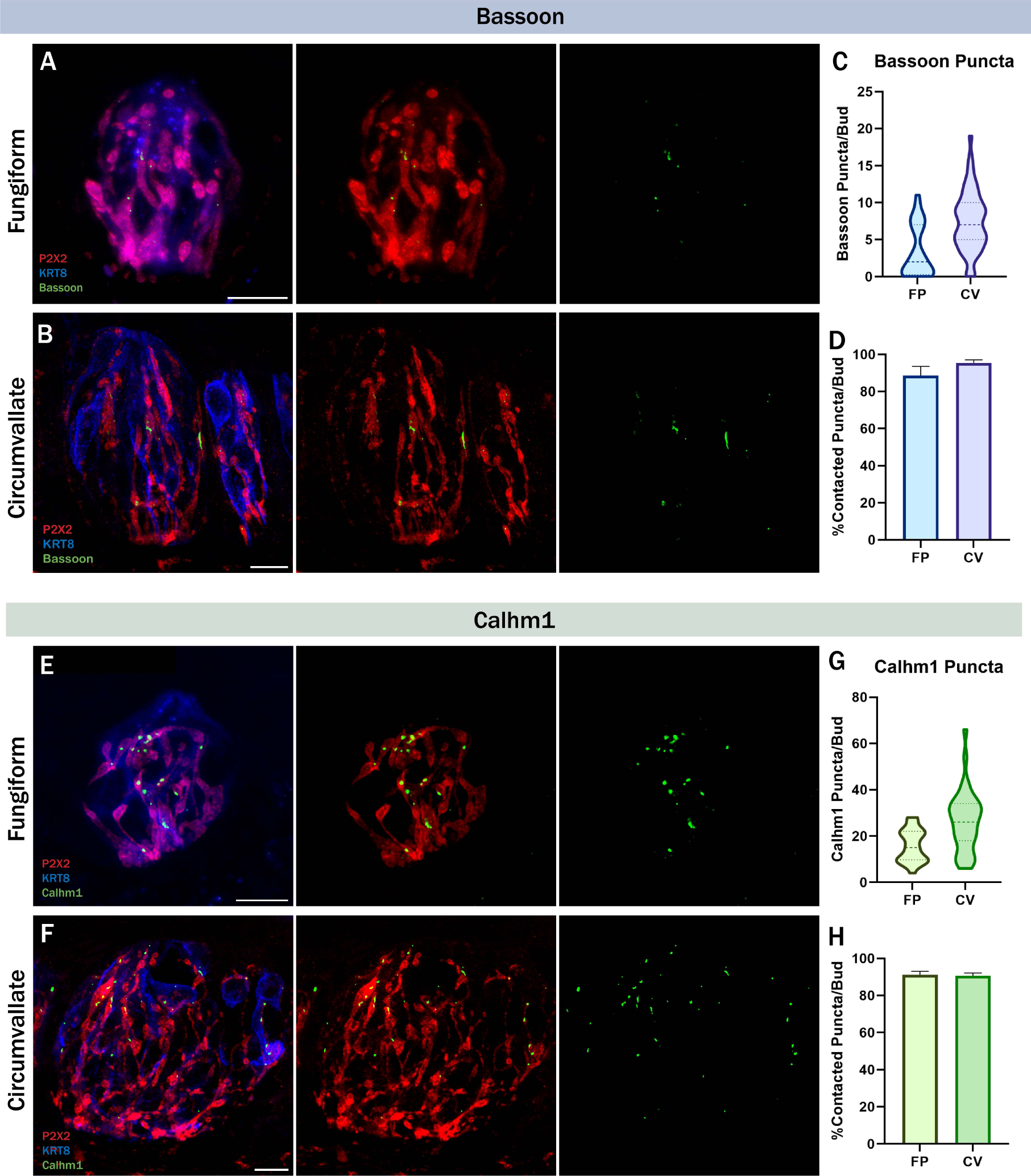
Bassoon and Calhm1 are frequently contacted by gustatory neurons. Immuno-labeling of P2X2 (red), KRT8 (blue), and Bassoon (green) in A) circumvallate and B) fungiform taste buds. C) Number of bassoon puncta per taste bud in fungiform and circumvallate taste buds. D) Average percentage of bassoon puncta that were contacted by P2X2-positive nerve fibers per taste bud. C&D) FP: N=36 buds/4 mice, CV: N=57 buds/4 mice. Immuno-labeling of P2X2 (red), KRT8 (blue), and CALHM1 (green) in E) circumvallate and F) fungiform taste buds. G) Number of CALHM1 puncta per taste bud in fungiform and circumvallate taste buds. H) Average percentage of CALHM1 puncta that were contacted by P2X2-positive nerve fibers per taste bud. G&H) FP: N=26 buds/4 mice, CV: N=39 buds/4 mice. Scale: 10µm. Error bars: SEM.

### Gustatory nerve transection leads to complete denervation of taste buds by day 4 and triggers TRC loss

To determine whether TRCs depend on gustatory neuron innervation to produce presynaptic sites, we first removed gustatory nerve fibers by nerve transection. Although the denervation of gustatory neurons from taste buds is essential for this study, it also has deleterious effects for the taste bud. Since gustatory nerve derived trophic factors, such as R-Spondin are necessary to stimulate taste stem cells to replenish the dying TRCs, nerve transection pauses TRC regeneration (Lin et al., 2021). As a result, transection of the chorda tympani (CT) nerve results in a substantial decrease in taste bud number, taste bud size, and taste cell numbers in fungiform papillae (Hendricks et al., 2002; Guagliardo and Hill, 2007; Li et al., 2015). Similarly, transection of the glossopharyngeal nerve results in almost complete loss of all circumvallate taste buds (Guth, 1957; John et al., 2003). Therefore, in order to analyze denervated taste buds with a minimal loss of TRCs, we needed to establish the earliest timepoint following nerve transection where afferent fibers have degenerated completely. To do this, we performed bilateral transection of the CT nerve to denervate fungiform taste papillae and bilateral transection of the glossopharyngeal nerve to denervate circumvallate taste papillae. We found that nerve transection results in a complete loss of afferent gustatory fibers within both fungiform and circumvallate taste buds by the 4^th^ day following surgery (Figure 2a,e). To assess the extent of TRC loss 4 days post nerve transection, immunostaining was conducted using Trpm5 and Car4 markers, which identify type II and type III cells, respectively. A significant loss of type II (Trpm5) cells in both fungiform (∼47%) and circumvallate (∼28%) taste buds was observed (Figure 2 c,g), which is expected due to their average lifespan of about 8 days (Perea-Martinez et al., 2013). While there was a loss of type III (Car4) cells, it was not significant in either fungiform (∼3%) and circumvallate (∼16%) taste buds (Figure 2d,h), which is also expected since they have a much longer lifespan of about 22 days (Perea-Martinez et al., 2013).

**Figure 2:**
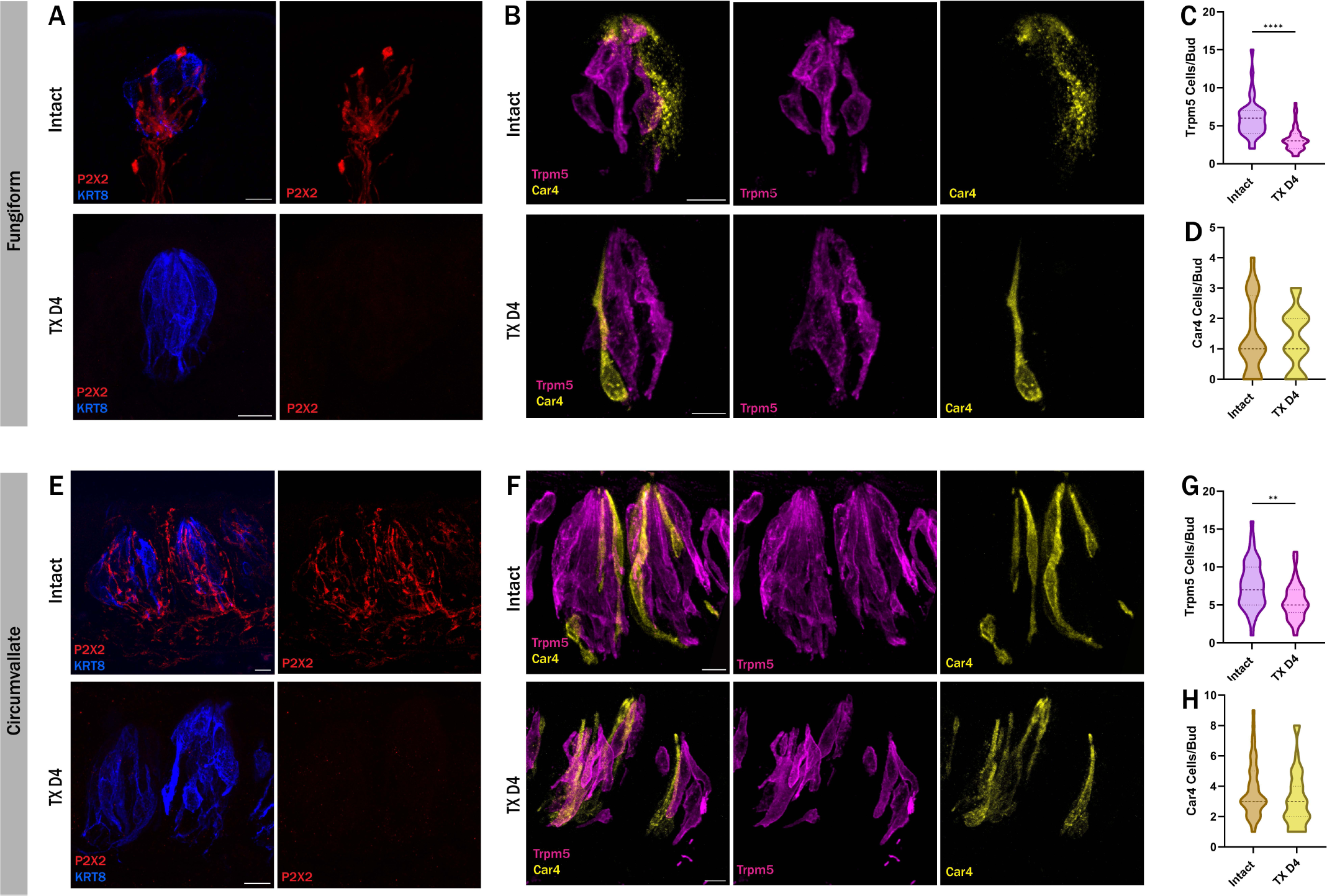
Nerve denervation induces loss of type II and type III taste cells. Comparison of immuno-labeled taste buds from intact control mice and denervated mice four days post nerve transection. A&E) Complete degeneration of nerve fibers occurs as early as four days following nerve transection in (A) fungiform and (E) circumvallate taste buds. This loss of innervation is accompanied by a loss in type II calls, marked by Trpm5 (magenta), and type III cells, marked by Car4 (yellow) in (B) fungiform and (F) circumvallate taste buds. Quantification of taste cells types in intact and transected mice reveals a significant loss of type II cells (C&G) while the loss of type III cells is not significantly different (D&H) in the transected groups. Intact FP: N=40buds/4mice, Transected FP: N=50buds/5mice, Intact CV: 45buds/3mice, Transected CV: 35buds/3mice. Scale: 10µm. Significance calculated using Mann-Whitney U: *P<0.0332, **P<0.0021, ***P<0.0002, ****P<0.0001.

### Gustatory nerve denervation results in a significant reduction of presynaptic sites

Because of the high rates of association between TRC presynaptic sites and gustatory neurons (∼90%), we hypothesize that gustatory innervation promotes or maintains presynaptic sites within the taste bud. Therefore, we predict that loss of gustatory innervation within the taste bud will reduce the number and/or intensity of Bassoon and CALHM1 immunolabeled puncta. To test this, we performed bilateral CT transections to eliminate fungiform innervation, and bilateral glossopharyngeal nerve transections to remove gustatory neurons from circumvallate taste buds. Strikingly, Bassoon staining was almost entirely abolished in fungiform and circumvallate taste buds 4 days following nerve transection compared to intact controls (Figure 3a-b, d-e). To account for taste cell loss following nerve transection, Bassoon puncta were normalized to the corresponding volume of mature TRCs per bud using Krt8 (cytokeratin 8) staining, which is a marker for all mature TRCs (Knapp et al., 1995; Mbiene and Roberts, 2003). This shows that the loss of Bassoon puncta is not a result of the reduction of mature taste cells due to normal TRC die-off in the absence of replenishment (Figure 3 c,f). Additionally, Type III cell loss, specifically, cannot account for the ablation of Bassoon puncta, as there are still 85-95% of Type III cells remaining 4 days following nerve transection (Figure 3d,h).

**Figure 3:**
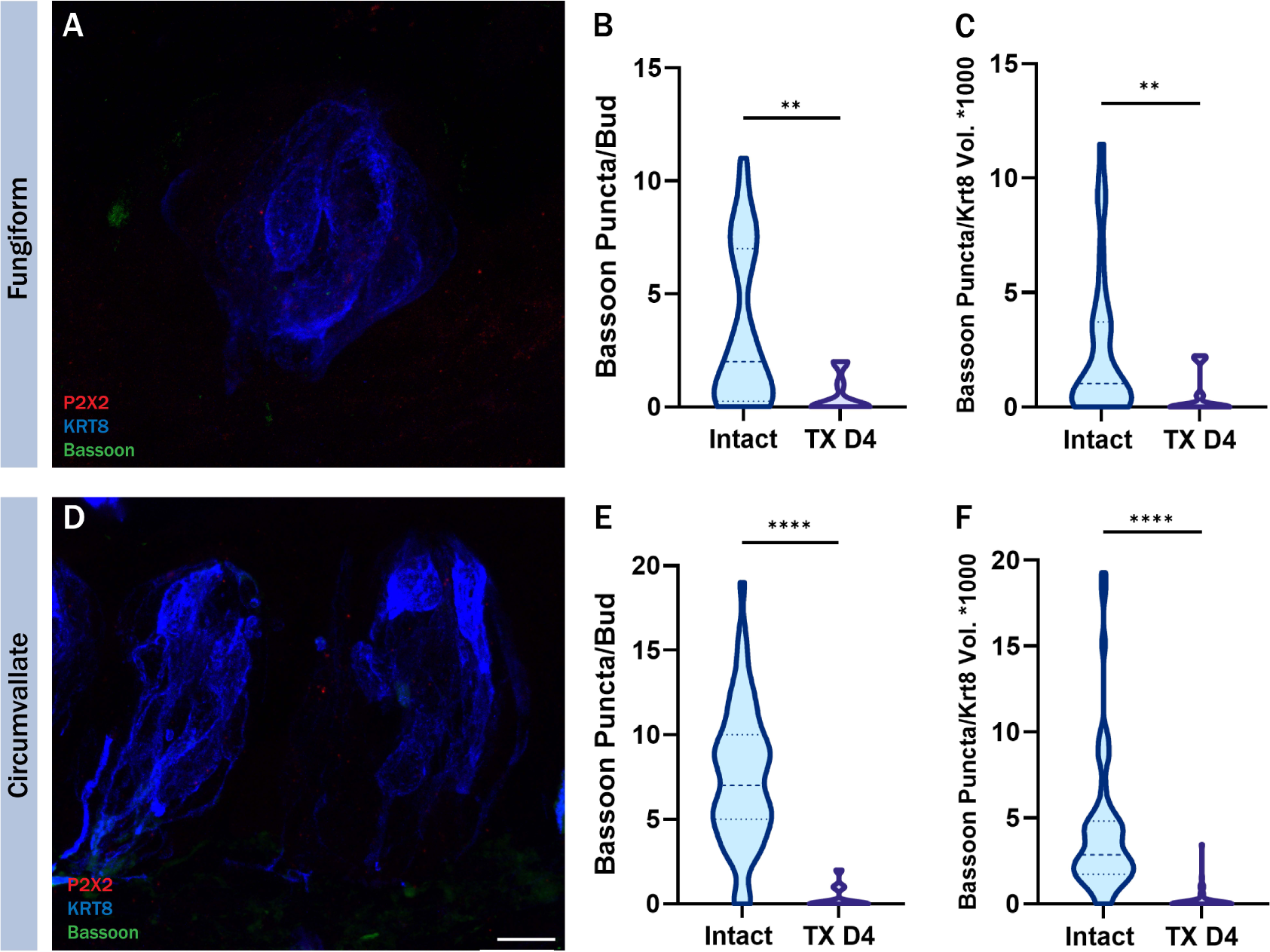
Bassoon puncta loss following nerve transection. Immuno-labeling of P2X2 (red), KRT8 (blue), and Bassoon (green) in transected day 4 (TX D4), (A) circumvallate and (D) fungiform taste buds. Number of Bassoon puncta per taste bud in intact and transected (B) fungiform and (E) circumvallate taste buds. Number of Bassoon puncta normalized to KRT8 volume per taste bud in intact and transected (C) fungiform and (F) circumvallate taste buds. CV: intact N=57 buds/4 mice, TX D4 N=61 buds/4 mice. FP: Intact N=36 buds/4 mice, TX D4 N=11 buds/2 mice. Scale: 10µm. Significance calculated using Mann-Whitney U: *P<0.0332, **P<0.0021, ***P<0.0002, ****P<0.0001.

Similarly, CALHM1 puncta are almost entirely lost following nerve transection. In fungiform papilla, CALHM1 puncta are almost completely gone 4 days post nerve transection (Figure 4a-b). This reduction is independent of mature TRC loss (Figure 4c) and cannot be attributed specifically to the loss of type II cell loss, given that they diminish about 45% by day four following nerve transection (Figure 2c). Circumvallate papillae, on the other hand, show a different pattern of CALHM1 puncta loss. CALHM1 loss was significant 4 days following transection compared to intact controls, however, a fair number of puncta persisted (Figure 4d,f). We extended our examination to taste buds 5 days following nerve transection and found that while more puncta are lost, some persist (Figure 3 e,f). CALHM1 puncta loss mirrored the loss of mature taste cells (Figure 4h,i) and type II cells (Figure 2g). Additional analysis reveals that even though total number of CALHM1 puncta diminishes slower in the circumvallate papillae, the intensity of remaining CALHM1 puncta is largely reduced (Figure 4g). Thus, CALHM1 is indeed severely impacted by neuronal denervation in both circumvallate and fungiform taste buds. Together, these findings reveal that normal accumulation of presynaptic machinery in TRCs is dependent on nerve fiber innervation.

**Figure 4:**
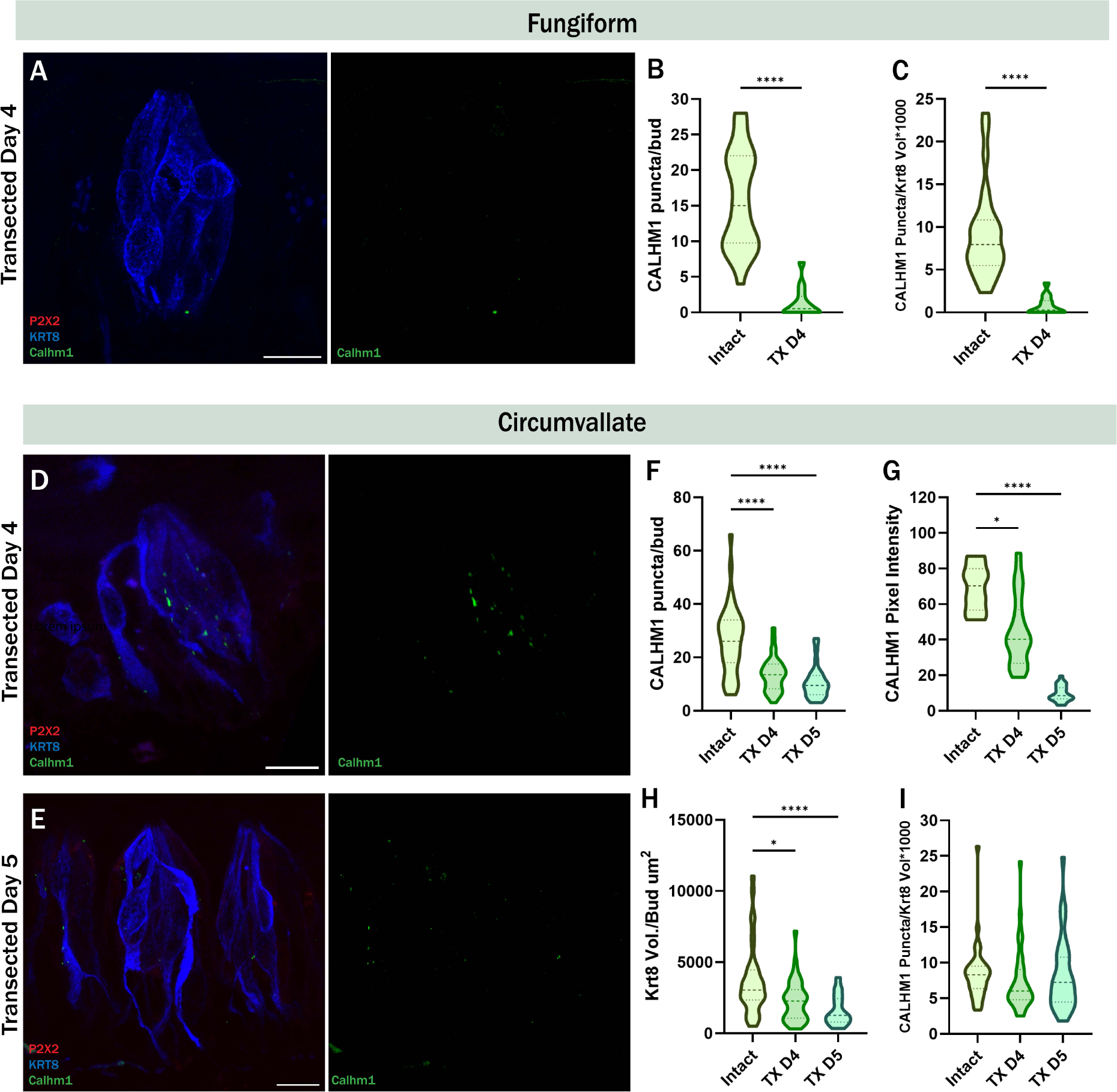
Bassoon puncta loss following nerve transection. Immuno-labeling of P2X2 (red), KRT8 (blue), and Bassoon (green) in transected day 4 (TX D4), (A) circumvallate and (D) fungiform taste buds. Number of Bassoon puncta per taste bud in intact and transected (B) fungiform and (E) circumvallate taste buds. Number of Bassoon puncta normalized to KRT8 volume per taste bud in intact and transected (C) fungiform and (F) circumvallate taste buds. CV: intact N=57 buds/4 mice, TX D4 N=61 buds/4 mice. FP: Intact N=36 buds/4 mice, TX D4 N=11 buds/2 mice. Scale: 10µm. Significance calculated using Mann-Whitney U: *P<0.0332, **P<0.0021, ***P<0.0002, ****P<0.0001.

### Gustatory denervation results in a reduction of CALHM1 and Bassoon mRNA expression

Loss of CALHM1 and Bassoon staining in de-innervated taste buds could be the result of transcriptional down regulation or internalization/dispersal of the proteins, within the taste cells. To test whether the transcription of Bassoon and CALHM1 is affected by gustatory innervation, we performed fluorescence in-situ hybridization (FISH) of Bassoon and CALHM1 transcripts in intact and day-4 transected CV papillae. We found that both Bassoon and CALHM1 transcripts are significantly reduced in CV tissues from mice 4 days after glossopharyngeal nerve transection compared to intact controls (Figure 5). The average loss of Bassoon transcript, (∼61.4%) far exceeds the expected loss due to reduction of type III cells (∼16.2%) 4 days following nerve transection (Figure 2h, Figure 5g). Similarly, the average loss of CALHM1 transcript (∼72.2%) is far more than the loss expected due to the average type II cell loss (∼28.3%) following nerve transection on day 4 (Figure 2g, Figure 5g). These data show that, without gustatory neuron innervation, the remaining taste cells produce less *Bassoon* and *CALHM1* mRNA transcripts which likely leads to the reduction of immunoreactivity in TRCs. However, we cannot rule out that internalization or the loss of localization by dispersal of protein from specific membrane sites may also occur.

**Figure 5:**
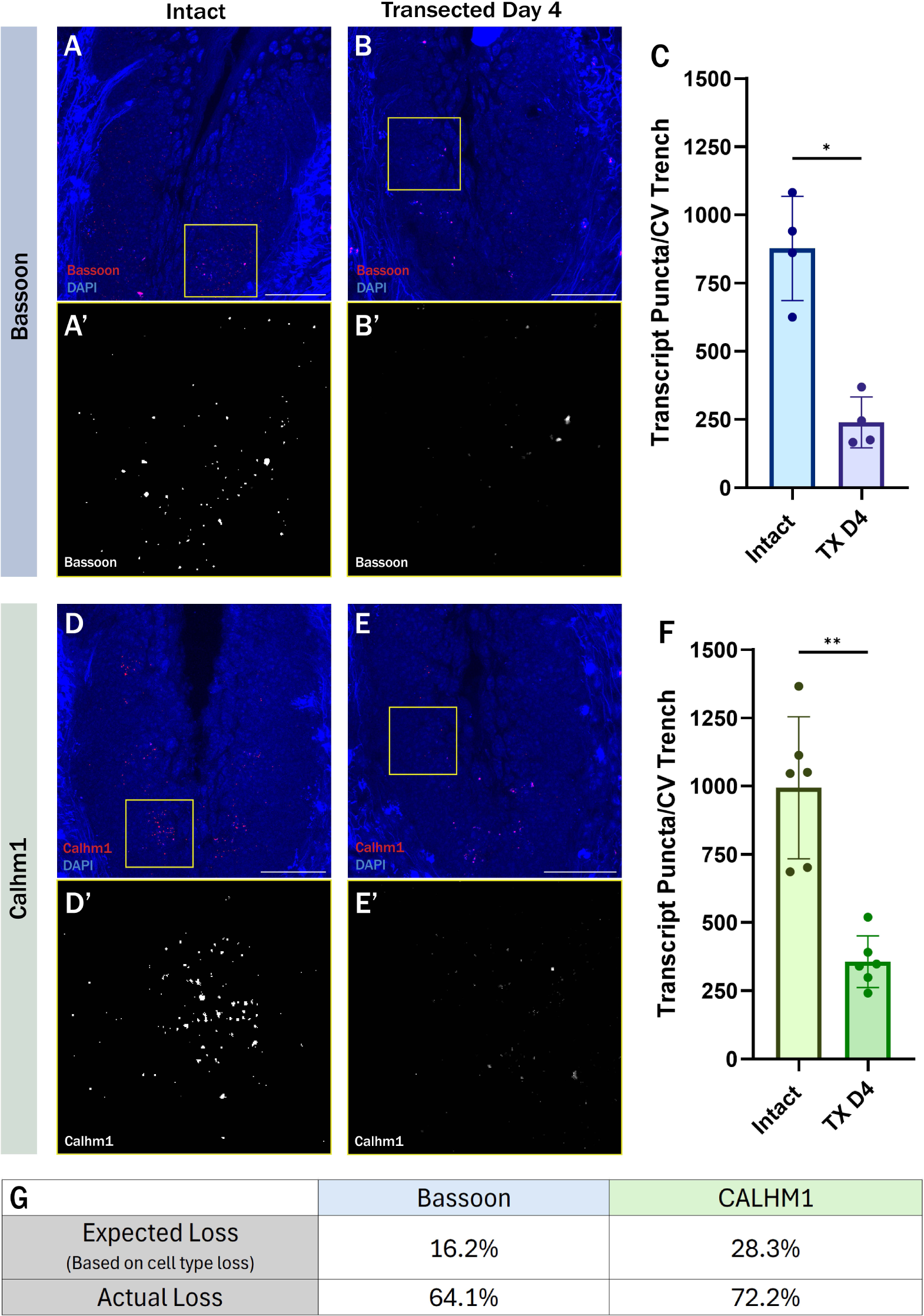
Bassoon and CALHM1 transcription are reduced following nerve transection. FISH labeling of Bassoon in (A) intact and (B) transected day 4 circumvallate papillae. A’&B’) Enlargement of yellow inset showing greyscale Bassoon labeling without DAPI. C) Bassoon transcripts quantified as puncta per trench in intact and transected circumvallate papillae. FISH labeling of CALHM1 in (D) intact and (E) transected day 4 circumvallate papillae. D’&E’) Enlargement of yellow inset showing greyscale CALHM1 labeling without DAPI. F) CALHM1 transcripts quantified as puncta per trench in intact and transected circumvallate papillae. G) Tab comparing the actual loss of transcripts vs the expected reduction based on type III cells (for Bassoon) and type II cell (for CALHM1) loss following transection. Bassoon: N=4 tranches/2 mice. Calhm1: N=6 trenches/2 mice. Scale: 50µm. Significance calculated using Mann-Whitney U: *P<0.0332, **P<0.0021, ***P<0.0002, ****P<0.0001. Error bars: Standard deviation.

### Gustatory neuron re-innervation following chorda tympani nerve transection

Early studies documenting the regeneration, innervation, and functional recovery of the CT nerve following injury in gerbils found that there was a delay between taste bud innervation and functional recovery of CT nerves (Cheal and Oakley, 1977; Cheal et al., 1977). Taste responses began to return in CT recordings about 12 days following nerve crush (Cheal et al., 1977), whereas nerve re-innervation was observed as early as 9 and 10 days following nerve crush (Cheal and Oakley, 1977). This delay could indicate that synaptic contact is not established immediately following nerve re-innervation into the taste bud. In mice, it has been shown that taste responses begin to reappear about 3 weeks following CT nerve crush. However, to our knowledge, there are no existing reports mapping the early timing of CT nerve re-innervation after injury. Therefore, we used in vivo and ex-vivo imaging techniques to observe nerve re-innervation following CT nerve injury.

To observe regenerating gustatory neurons, we monitored nerve regrowth in individual fungiform taste buds using intravital 2-photon imaging of Phox2b-cre;TdTomato mice who had undergone unilateral CT transection. In these animals, all gustatory neurons are marked with bright red fluorescence. Images of the same taste buds were captured before and 14 days following unilateral nerve transection. Following the nerve cut, nerve fibers had largely dissipated by day 4, but began to regenerate and re-innervate taste buds by day 8 (Figure 6a-e). Interestingly, we found that nerve fiber volume began to return to normal levels by day 11 and showed even higher volumes by day 14. Upon further analysis, it appears that the rapid return of gustatory neurons may not be located within the taste bud area. The sampling area for volume analysis was taken from the whole fungiform papillae rather than from the taste bud area only. On days 11 and 14, the nerve fibers appear to be more spread out compared to the intact side (Figure 6b,c), which may be due to more fibers innervating the perigemmal space (the area directly adjacent to a taste bud) rather than within the actual taste bud. To corroborate the 2-photon analysis, we immunostained fungiform taste buds and saw that gustatory neuron re-innervation does indeed begin to occur approximately 8 days following nerve transection (Figure 6F). However, nerve fiber volume within the taste bud is still significantly less than intact controls on days 12 and 14 (Figure 6F). When the location of re-innervating gustatory neurons is observed, many gustatory neuron branches are seen innervating the perigemmal space on day 12 (Figure 6). So while the total volume of gustatory neurons within fungiform papillae returns by day 12 (figure 6a-e), the portion that are found innervating the taste bud remains less than intact controls by day 12 (Figure 6 f,g). The data shown here is consistent with previous reports showing that gustatory neuron volume within the taste bud does not reach normal levels until 8 weeks or longer following CT nerve injury (Meng et al., 2017; Dong et al., 2023).

**Figure 6:**
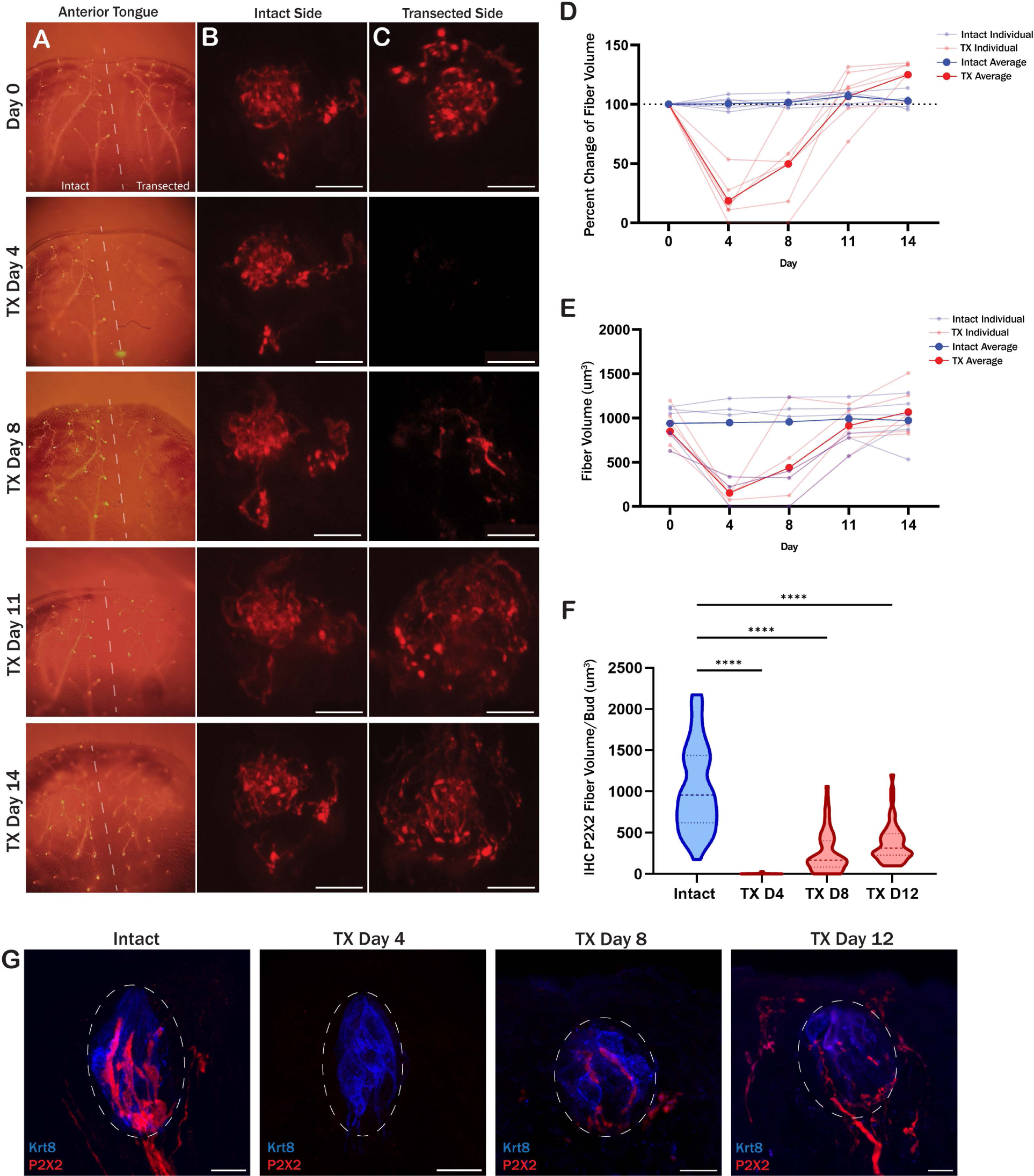
Regrowth of the chorda tympani nerve into fungiform papilla following nerve transection. Gustatory nerve regrowth was tracked in fungiform papilla, in-vivo and ex-vivo, following CT nerve transection. A-E) Phox2b-cre;TdTomato animals who underwent unilateral CT nerve transection were used for intravital imaging of the anterior tongue and fungiform papillae to visualize gustatory nerve innervation. A) Epifluorescent images of the anterior portion of the tongue over the course of 14 days, with a dashed line splitting the intact left side and the CT transected right side. 2-photon images of the same taste bud imaged over 14 days on the (B) intact side and (C) transected side of the togue. D&E) Nerve fiber volume was quantified in intact (blue) and transected (red) taste buds over time and is shown as (D) percent change from day-0 and (E) absolute nerve fiber volume. F) Gustatory marker, P2X2, was quantified within the taste bud via immunostaining in intact and bilateral CT nerve transected mice 4-, 8-, and 12-days following surgery (50-60 buds/condition). G) Gustatory neuron (P2X2) volume from intact and transected taste buds was quantified only from the taste bud area positive for Krt8 staining, indicated with white dashed circles. Scale: 10 µm. Significance calculated using Kruskal-Wallis: ****P<0.0001.

### Chorda Tympani nerve re-innervation stimulates the return of Bassoon and CALHM1 independent of nerve fiber contact

To assess the effect of nerve re-innervation on Bassoon and CALHM1 accumulation in TRCs, we collected tissues from intact, transected day-4, -8, and -12 mice for immunostaining to determine when synapses are established (Figure 7). As shown earlier, Bassoon and CALHM1 puncta were almost entirely abolished on the 4th day following nerve transection (Figure 7 e,k). On day 8, when nerve fibers just start to return to the taste bud, Bassoon and CALHM1 also began to come back (Figure 7 e,k). By day 12, average Bassoon puncta per bud exceeds that of intact controls (Figure 7 d,e), while CALHM1 puncta had not yet returned to normal levels (Figure 7 j,k). To determine whether nerve fiber contact is necessary for Bassoon and CALHM1 accumulation in taste cells, we quantified the percentage of Bassoon and CALHM1 puncta that are contacted by P2X2 nerve fibers during nerve re-innervation. When nerve fibers begin to re-innervate the taste bud 8 days post nerve transection, we found that Bassoon and CALHM1 puncta are contacted by P2X2 nerve fibers significantly less than intact controls (Figure 7 f,l). Nerve fiber contact does not return to near-normal levels until 12 days following nerve transection (Figure 7 f,l). These findings illustrate how the accumulation of Bassoon and CALHM1 depends on the presence of nearby nerve fiber innervation, but not direct nerve contact. If nerve fiber contact was indeed necessary for taste cells to produce Bassoon and CALHM1, the rate of contacted Bassoon and CALHM1 puncta would remain at a high level, even at the early stages of re-innervation.

**Figure 7:**
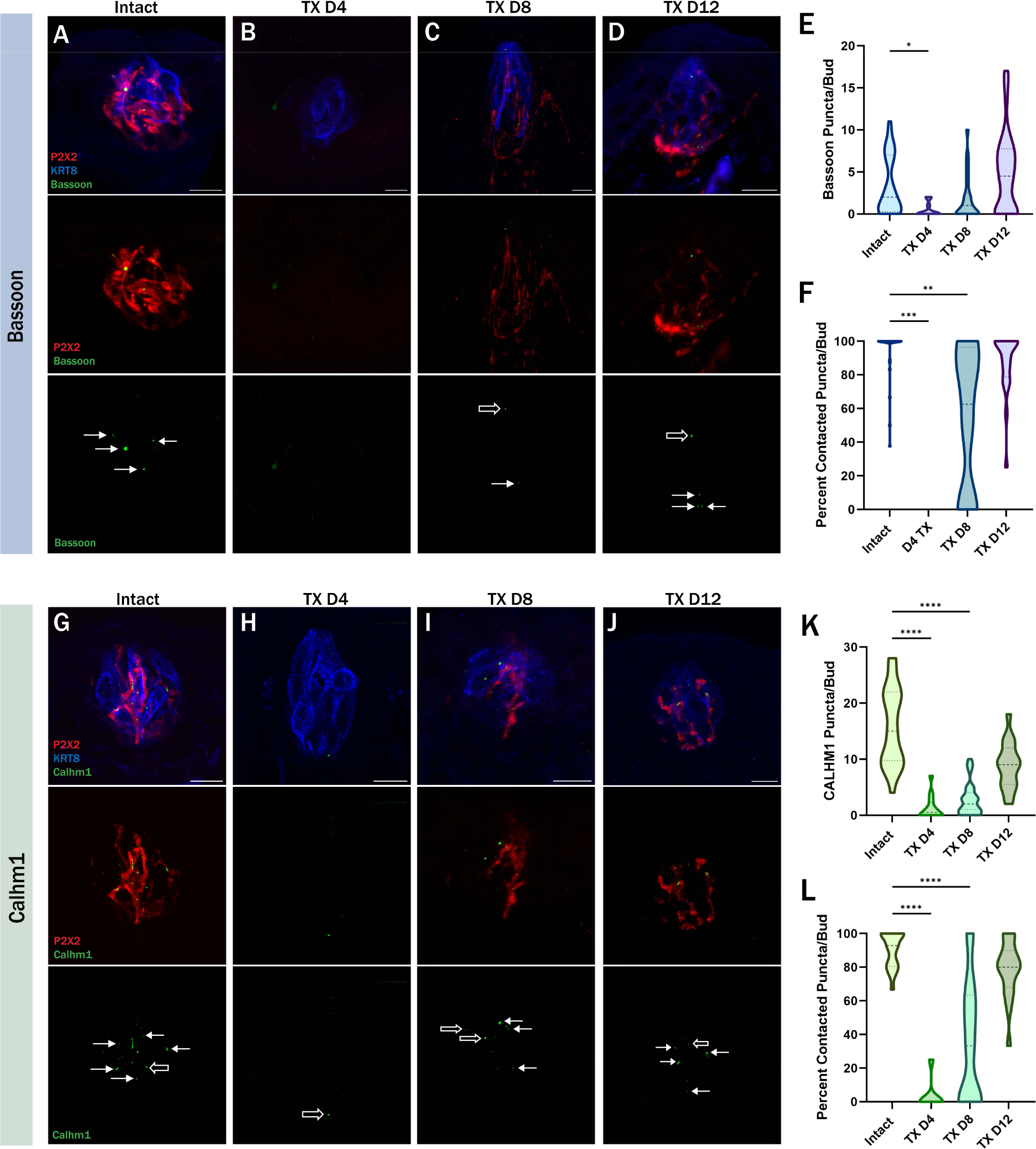
Fewer Bassoon and CALHM1 puncta are contacted by gustatory neurons in the early stages of re-innervation than in intact taste buds. Immuno-labeling of P2X2 (red), KRT8 (blue), and Bassoon (green) in (A) intact, (B) transected day 4, (C) transected day 8, and (D) transected day 12 fungiform taste buds. E) Number of Bassoon puncta per taste bud. F) Average percentage of Bassoon puncta that are contacted by P2X2-positive nerve fibers per taste bud. E&F) Intact: N=36 buds/4 mice, TX D4: N=11 buds/2 mice, TX D8: N=25 buds/3 mice, TX D12: N=32 buds/4 mice. Immuno-labeling of P2X2 (red), KRT8 (blue), and CALHM1 (green) in (G) intact, (H) transected day 4, (I) transected day 8, and (J) transected day 12 fungiform taste buds. K) Number of CALHM1 puncta per taste bud. L)Average percentage of CALHM1 puncta that are contacted by P2X2-positive nerve fibers per taste bud. K&L) Intact: N=26 buds/4 mice, TX D4: N=14 buds/2 mice, TX D8: N=32 buds/4 mice, TX D12: N=17 buds/4 mice. Scale: 10µm. Significance calculated using Kruskal-Wallis: *P<0.0332, **P<0.0021, ***P<0.0002, ****P<0.0001.

## Discussion

The data we reveal here unveils novel insights into the intricate mechanisms governing synaptogenesis and synaptic plasticity within the dynamic and ever-evolving environment of taste buds. By associating TRC presynaptic sites with gustatory nerve endings, we reveal initial mechanisms of synapse formation within the taste bud. Under normal conditions, presynaptic sited (marked by Bassoon and CALHM1) are present and frequently contacted by gustatory nerve fibers. However, transection of the chorda tympani and glossopharyngeal nerves causes a significant decrease in Bassoon and CALHM1 puncta, the extent of which cannot be explained solely by the loss of TRCs in denervated taste buds. We found that mRNA transcripts for Bassoon and CALHM1 are both reduced after nerve transection, suggesting that the presence of nerve fibers stimulates the production of Bassoon and CALHM1. However, we cannot rule out the possibility of additional changes to protein localization or degradation. Re-innervation experiments reveal that as the CT nerve regrows, Bassoon and CALHM1 puncta begin to reappear in the taste buds, though they are not contacted as frequently by the newly innervated nerve fibers. This indicates that the initial stages of nerve fiber regrowth can stimulate taste cells to produce presynaptic sites independently of direct nerve contact. We hypothesize that there are neuronally derived trophic factors (perhaps R-Spondin [see below discussion]) that prompt taste cells to form presynaptic sites; however, further research is necessary to substantiate this theory.

Although presynaptic markers have been an excellent tool to observe synapse formation, using these presynaptic markers alone cannot precisely map synapse formation in specific cell types. Analysis of 3D electron micrographs has shown hybrid cells with both vesicular and channel synaptic contacts, though, the identity of these cells are not yet known (Wilson et al., 2021). Therefore, there are some cells that may have both channel synapses using the CALHM1/3 channels and also vesicular synapses marked by Bassoon. The addition of taste cell type markers can show how frequently this occurs; however, we were not able to accomplish this due to a lack of antibody compatibility.

Studies mapping synapse turnover in the brain are able to document synapse plasticity by observing the addition and removal of dendritic spines over time (Attardo et al., 2015), as gross axonal and dendritic structures are mostly stable in the adult brain (Südhof, 2021). However, due to the dynamic nature of the taste bud, the peripheral endings of gustatory neurons are highly plastic. Anatomical analysis describes gustatory nerve endings taking on diverse morphologies, such as branch retraction and sprouting, indicating states of plasticity (Ohman et al., 2023). Moreover, intravital studies reveal that gustatory neurons exhibit remarkable plasticity (Whiddon et al., 2023; Wood et al., 2024), with peripheral arbors averaging a branch extension or retraction every 8 hours (Whiddon et al., 2023). Because of this rapid plasticity, the ability to observe synapse turnover in the taste bud remains a challenge. Therefore, implementing a genetic synapse labeling system like GRASP (GFP Reconstitution Across Synaptic Partners) could be an effective tool to track synapse turnover in taste buds over time. GRASP labels synapses using endogenous fluorescence (Kim et al., 2012; Macpherson et al., 2015), that could, theoretically, enable the observation of synapse formation and dissolution within the taste bud. Data from such experiments would provide a comprehensive understanding of synapse turnover in taste buds.

Taste bud homeostasis depends on a complex interplay between taste receptor cells and gustatory neurons. In fact, when nerve fiber innervation is interrupted, taste buds slowly deteriorate because taste cell turnover is interrupted. TRCs can only replace themselves when there is a steady supply of neuronally derived R-Spondin. The Wnt pathway in taste stem cells is positively regulated by R-Spondin through LGR4/5 receptors. Thus, neuronal supply of R-Spondin stimulates the replacement of old taste cells. It is tempting to hypothesize that R-Spondin may also stimulate taste receptor cells (TRCs) to produce presynaptic sites. If this hypothesis holds true, administering exogenous R-Spondin after nerve transection could potentially rescue the loss of presynaptic sites in taste cells. However, if R-Spondin is not the molecular trigger for presynaptic site production, this experiment could be repeated using any other putative trophic factor identified through RNA-seq experiments.

Well-studied central networks have not yet worked out basic processes that are integral to synaptogenesis (Südhof, 2021). Synapse formation is traditionally thought to be orchestrated first by axon guidance mechanisms, positioning the axon adjacent to its target, then synaptic adhesion molecule (SAM) compatibility determines which neurons form synapses at what location (Südhof, 2021). These trans-synaptic adhesion molecules dictate the formation and dictate activity-dependent function of synapses (Kim et al., 2021). There are many known SAMs postulated to initiate synapse formation, though, none are documented in taste cells or gustatory neurons to interact in the trans-synaptic space. Gustatory neurons express Gaba receptor a1 (Dvoryanchikov et al., 2011, 2017), a SAM that has been shown to interact with neurexins on the presynaptic cell within the hippocampus (Panzanelli et al., 2017). Additionally, sweet and bitter TRCs express semaphorin7a and - 3a, respectively, to guide gustatory neurons to make appropriate and specific connections (Lee et al., 2017), while protocadherin-20 has been proposed to guide gustatory neurons to connect specifically to sweet and umami cells (Hirose et al., 2020). These molecular tags may help guide gustatory neurons to specific taste cells, though, though exact mechanisms governing the formation of synaptic connections is entirely unknown. However, the findings described here, have illuminated the dynamic relationship between nerve fibers and presynaptic sites within taste buds, demonstrating that nerve innervation, rather than contact, directly influences the presence and maintenance of synapse-related proteins in TRCs. This fundamental mechanism provides a solid foundation for future studies investigating synapse formation within taste buds.

## Methods

### Animals

All procedures were conducted in accordance with US National Institutes of Health (NIH) guidelines for the care and use of laboratory animals and were approved by the University of Texas at San Antonio IACUC. Both male and female mice were used in this study. For immunohistochemical experiments of sectioned tongue tissues, C57/b6 mice (Jax Strain #000664) were used along with Phox2b-Cre;tdTomato double transgenic mice (Jax Strains #016223, #007909). All mice were between 3 and 8 months old; weighing between 18 and 35g. Food and water were available ad libitum. The mouse colony was maintained on a regular 12/12 h light-dark cycle. Animal numbers are reported in the figure legends.

### Nerve transection surgery

Aseptic bilateral nerve sectioning of the glossopharyngeal nerve and the chorda tympani nerve were performed under aerosolized isoflurane anesthesia at 5% in oxygen (4L/min) for induction and 1-3% in oxygen (0.5L/min) for maintenance. Body temperature was maintained with a heating pad. The CT nerve was approached ventrally in the neck and transected after it bifurcates from the lingual nerve with severed ends left in place as previously described (McCluskey, 2004; Padalhin et al., 2022). For the glossopharyngeal nerve transection, the nerve was accessed ventrally through the neck and visualized by retracting the omohyoid and sternoid muscles medially and the digastric muscle and sublingual/submandibular salivary glands laterally. The glossopharyngeal nerve was located inferiorly to the hypoglossal nerve and lateral to the pharyngeal branch of the vagus nerve, where it was then transected (John et al., 2003) The wound was closed with discontinuous sutures. Meloxicam ER and buprenorphine ER analgesics were given prior to surgery and animal well-being was monitored in the days following surgery.

### Immunohistochemistry

Mice were euthanized through CO2 inhalation followed by trans-cardial perfusion with 20mL of 1x PBS and 10mL of 4% paraformaldehyde solution in 1x M PBS, sequentially. Complete perfusion is critical for success of Bassoon and CALHM1 stains. Tongues were extracted and cryoprotected in 30% sucrose at 4 °C overnight. The tongues were then dissected to separate the circumvallate papillae (CV) and the anterior 2/3 of the tongue where the fungiform papillae are located. The anterior 2/3 of the tongue was then cut sagittally to split it into left and right halves. The left and right tongue tip and the CV were embedded in OCT compound and stored at -80 °C. Samples were cryo-sectioned at 18 μm thickness and mounted directly onto anti-frost slides. Sections of the CV and tongue tips underwent antigen retrieval by incubating in sodium citrate buffer (10mM Sodium Citrate, 0.05% Tween 20, pH:6) at 80°C for 20 minutes. The slides were removed from the sodium citrate buffer and allowed to cool to room temperature before washing with 1x PBS for 5 minutes. The slides were then dried along the edges before applying a hydrophobic barrier and dried for 5 more minutes. Blocking buffer was then applied to the sections: first, 5% donkey serum with 0.3% triton-x100 in 1x PBS at 25°C for 30 minutes, then blocked again with M.O.M.® (Mouse on Mouse) Blocking Reagent (MKB-2213-1) diluted to 1 drop in 1mL 1x PBS at 25°C for 30 minutes. Primary antibodies (Table 1) were diluted in 5% donkey serum with 0.3% triton-x100 in 1x PBS overnight at 4 °C. After 3 × 5-min washes with 1x PBS with 0.1% Trinto-x100, secondary antibodies (Table 1) were diluted to 1:1000 in 5% donkey serum with 0.3% triton-x100 in 1x PBS and incubated for 4 hr at 4 °C. The slides were then washed for 3 × 5 min with 1x PBS with 0.1% triton-x100 and then mounted with for imaging.

**Table 1:**
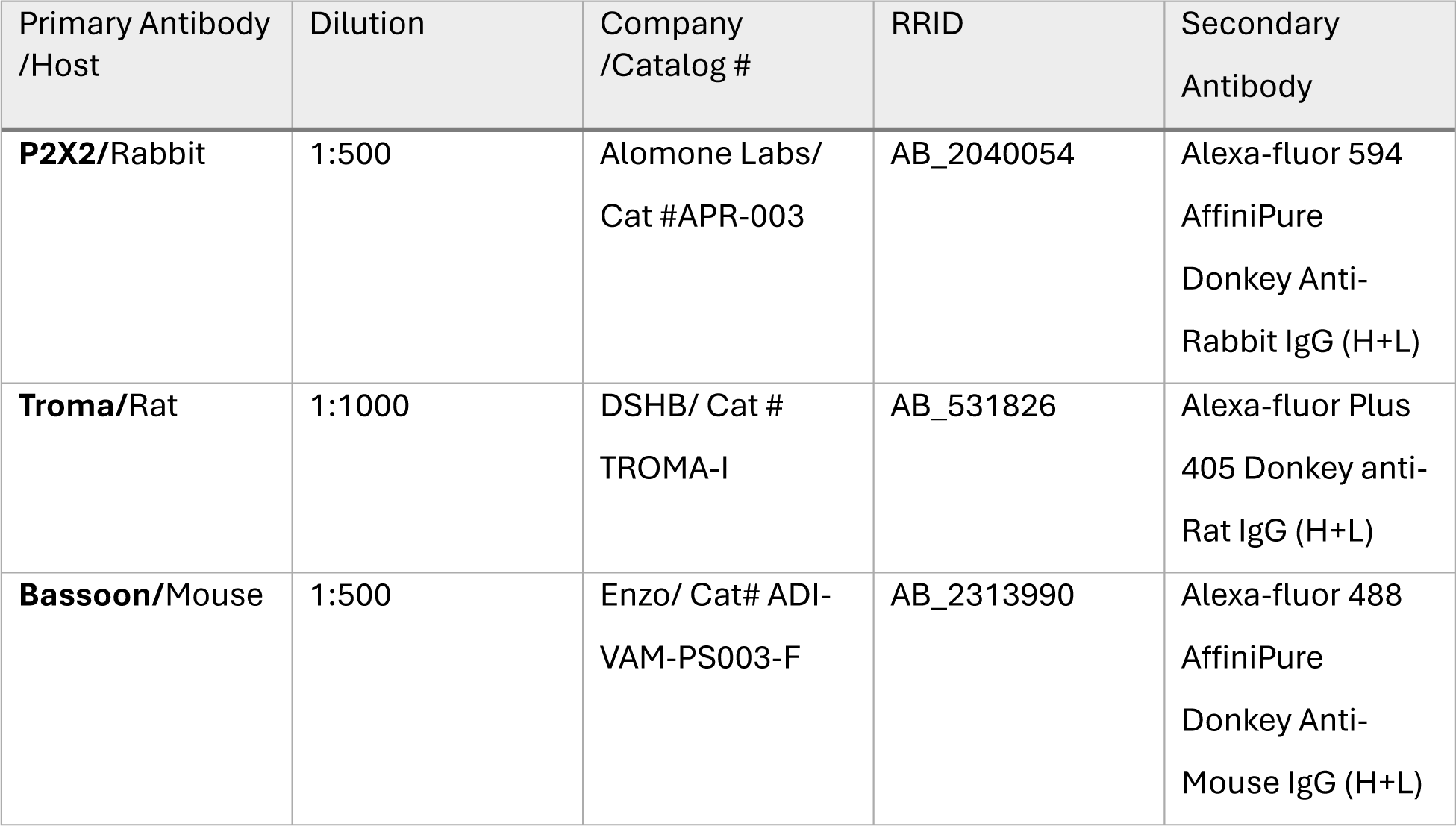
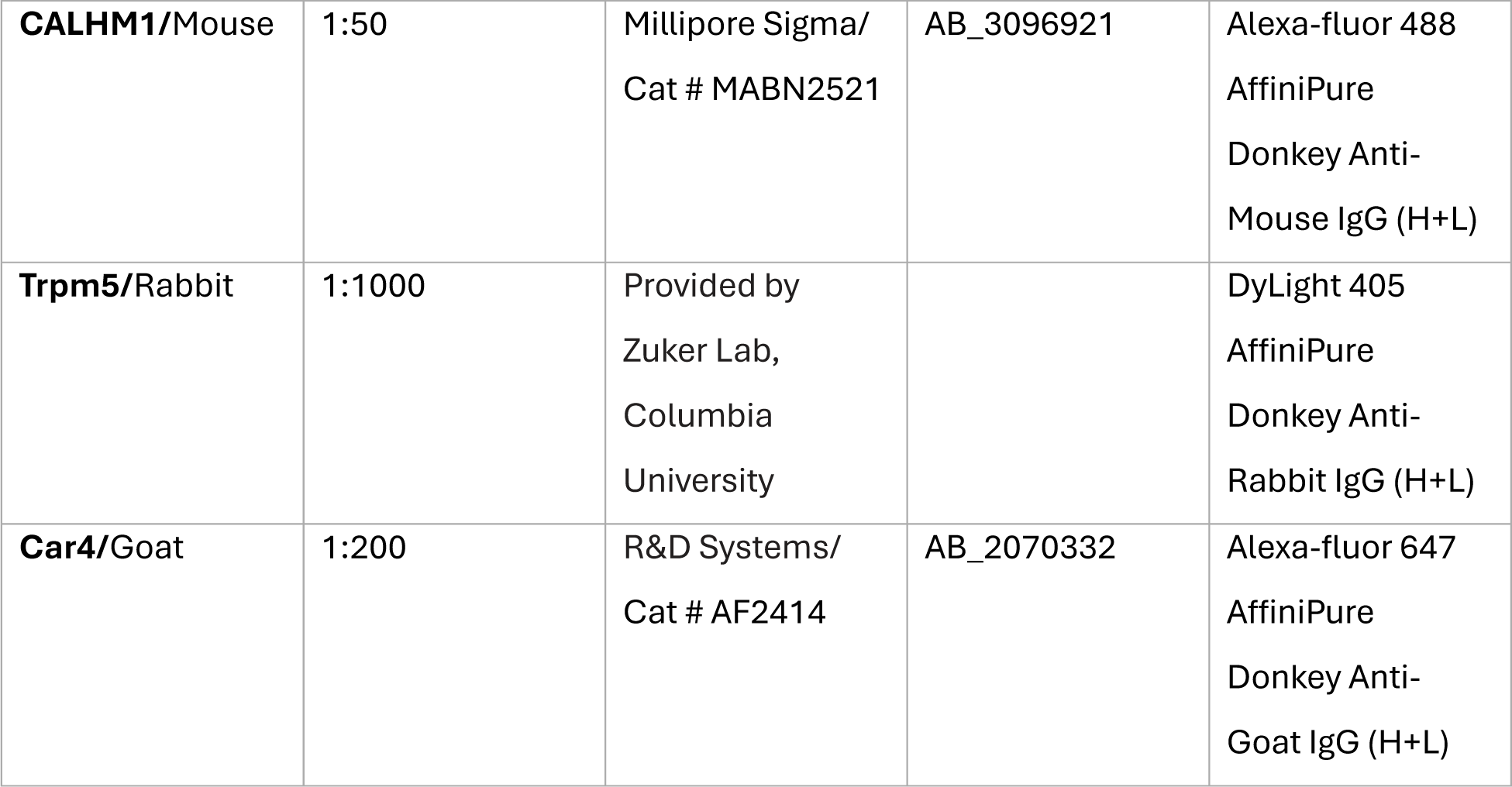
Primary and secondary antibodies used for immunohistochemistry of taste tissue.

### In situ hybridization

Mice were euthanized through CO2 inhalation and cervical dislocation. The CV of the tongue was quickly dissected, placed in ice-cold OCT medium, and flash frozen in dry ice. Frozen tissues were stored in -80 °C until ready to process. Samples were cryo-sectioned at 15 μm thickness and mounted directly onto anti-frost slides. The protocol for Manual RNAscope® Assay by ACD was followed using RNAscope® Fluorescent Multiplex Detection Kit (Catalog No. 320293). The probes used were RNAscope™ Probe-Mm-CALHM1 (Cat No. 558951) and RNAscope™ Probe-Mm-Bsn-C1 (Cat No. 1119681-C1). Following probe and amplifier hybridization, DAPI was applied to the slides for 30 seconds. Stained slides were then mounted with coverslips using VECTASHIELD® Antifade Mounting Medium (H-1000-10) and sealed with clear nail polish.

### Confocal image acquisition and quantification

A Zeiss 710 confocal microscope was used with the 63x objective to image the tongue sections for individual taste buds. Whole CV trenches for FISH staining were imaged using the 20x objective. Z-stack images consisted of 20-35 stacks at 0.6 μm steps. ImageJ (FIJI) was used for all immune-stained tissues. Taste bud area profile ROIs were determined by KRT8 staining and outlined using the lasso tool. To measure taste cell and gustatory nerve fiber volume for each taste bud, the z-stack fluorescence was adjusted to a median brightness with a 1.0 pixel radius and a threshold was set to encompass all positive signal. The same ROI used to determine bud area was used to measure volume. Using a macro (https://visikol.com/blog/2018/11/29/blog-post-loading-and-measurement-of-volumes-in-3d-confocal-image-stacks-with-imagej/), the thresholded z-stack was converted from voxels to μm^3^. Counting CALHM1 and Bassoon puncta was accomplished by using the magic wand tracing tool in ImageJ. CALHM1 and Bassoon puncta greater than 0.5 μm and small puncta clusters were counted as 1 individual puncta. Puncta contacts were evaluated by P2X2 staining. Puncta that were overlapping with or directly touching P2X2 staining were categorized as “contacted puncta” and all others were characterized as “uncontacted”.

Imaris image analysis software was used for in situ hybridization RNAscope images. Number of CALHM1 and Bassoon puncta were counted by using the surfaces analysis module (surface gain size: 0.300 μm, Manual threshold value: 1500, surface volume 0.100 μm^3^).

### Anesthesia and preparation of the mouse for imaging

Mice were given an initial IP injection of 10 mg/kg ketamine and 0.1 mg/kg xylazine (Covetrus). An IP injection was also given prior to positioning the mouse in the imaging apparatus to maintain fluid levels during imaging. The mouse was placed on his/her back and the tongue was inserted in the opening of a tongue holder using blunt forceps, as in (Wood et. Al. 2024). Initially, a 10× Olympus lens was used to spatially recognize that the same taste bud was being imaged. Once the correct bud was identified, a 40× water immersion lens was used to image the gustatory fibers. After each imaging session, the mouse was given an IP injection of Atipamezole (Covetrus) (1mg/kg) to reverse the effects of the sedative.

### Two-photon imaging

Two-photon excitation microscopy imaging was used to obtain 3-D images of a live mouse tongue, a Bruker system with Prairie View software. A laser strength of approximately 400 mV and a wavelength of 1100 nm was used to visualize the tdTomato-labeled nerve fibers innervating the tongue. The same PMT and imaging strength were used each day to ensure no variation due to the setting changes. Images were collected at 1024 × 1024 pixels, resonance galvo, with resonance averaging at 2 frames. This resonance setting was used because it takes a faster image which is critical due to our time constraints of the sedative. Z stacks were collected at 5 μm intervals covering approximately a volume of 100 μm, cubed. Typically, most of the taste bud is imaged within the first 50 μm of the Z-axis, but additional images were collected toward the base of the bud to help orient the 3-d volume as well as

### Three-dimensional image analysis

ImageJ and Imaris were utilized for data analysis and 3-D reconstruction of 2-photon images. ImageJ software was used to obtain a volume measurement of the fibers in pixels, cubed. First, a consistent threshold setting was used in ImageJ to minimize background fluorescence/autofluorescence. Then, ImageJ was used to extract the region of interest and remove extraneous pieces that were not included in the analysis. This region of interest included the bottom portion of the bud that comprises the dense papilla core, excluding the gustatory fibers beyond the basal lamina. A volume calculator in ImageJ was utilized to calculate the volume of fibers innervating each taste bud from each section of the z-stack. Imaris software was used to create 3-D reconstructions of the innervating fibers after ImageJ was used to select the region of interest. The “surface volume” setting was applied to the z-stack. This setting finds the edges of 3-D images and renders an outline of the 3-D image.

---

**Supplemental Figure 1:**
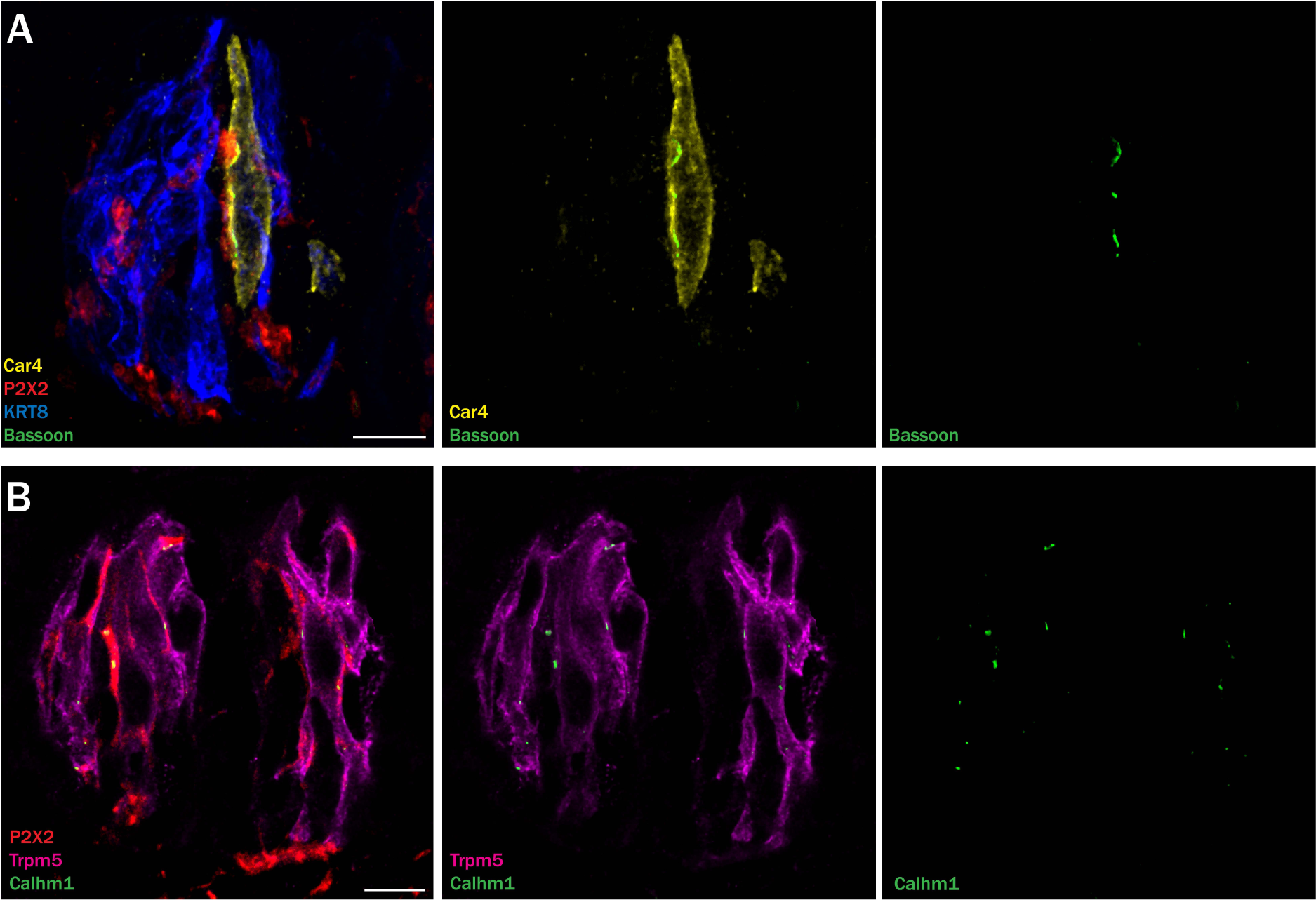
Bassoon and CALHM1 puncta colocalize with type III and type II cells, respectively. A) Immuno-labeling of Krt8 (blue), P2X2 (red), Car4 (yellow) and Bassoon (green). Bassoon staining is only se at the junction of Car4 cells and P2X2 neurons. Bassoon staining is only seen at the junction of Car4 (type III) cells and P2X2 neurons. B) Immuno-labeling of Trpm5 (Magenta), P2X2 (red), and CALHM1 (green). CALHM1 staining is only seen at the junction of Trpm5 (type II) cells and P2X2 neurons. Scale: 10µm

